# Is the Bitter Lesson coming to genomic prediction? An untuned tabular foundation model matches or beats specialized methods

**DOI:** 10.64898/2026.11.03.764067

**Authors:** Lex Flagel

## Abstract

Genomic prediction (GP) is dominated by models designed around assumptions about how genetic variation affects phenotypes. Here we asked how well a general-purpose pretrained tabular foundation model called TabPFN-3.5 does with no genetics-specific modeling and no tuning when given marker genotypes directly. We benchmarked TabPFN-3.5 using a published comparison of 18 GP methods on 18 traits in six plant species. TabPFN-3.5 ranked first on 11 of 18 traits and had the best median rank among all methods. Because the published benchmark did not identify its test sets, we reran GBLUP, RKHS and elastic-net models on the same splits as TabPFN-3.5. Against the best of these, TabPFN-3.5 was ahead on 12 of 18 traits, with significant gains on all three soybean traits and rice flowering time. In the soybean panel, which had the largest number of lines, its advantage was clearest. In leave-families-out cross-validation, where models were trained on some soybean families and used to predict lines from other families withheld from training, every model lost accuracy and TabPFN-3.5 performed on par with GBLUP and RKHS. Our results show that a general-purpose learner matches or exceeds decades of specialized GP models. This finding is consistent with Sutton’s “Bitter Lesson”, which holds that methods that scale with computation eventually overtake those built on human knowledge of the domain.

## 1 Introduction

Genomic prediction (GP) predicts an individual’s breeding value or phenotype from genome-wide markers (Meuwissen et al., 2001). A model is trained on individuals with both genotypes and phenotypes and then applied to individuals that have only been genotyped. Because genotyping is fast and cheap relative to phenotyping, GP lets breeders screen and select candidates before phenotypes exist. It has become routine in dairy cattle and in plant breeding programs across major crops (Hayes et al., 2009; Wiggans et al., 2017; Heffner et al., 2009; Crossa et al., 2017). As an example of the value of GP, the USDA estimated that genomic selection returned about $4 billion to the US dairy industry in its first nine years (Rexroad et al., 2019).

The rate of genetic gain scales linearly with prediction accuracy (Falconer and Mackay, 1996). Small gains in prediction accuracy therefore accumulate across generations, which has motivated two decades of GP method development. Formally, a GP dataset is an *N* × *M* matrix of marker genotypes (*N* individuals and *M* markers), an *N* × *P* matrix of phenotypes, and sometimes covariates. In machine learning parlance, GP is a standard supervised regression problem where we learn a mapping from a row of genotypes to a phenotype on individuals that have both, then apply it to individuals that only have genotypes.

Many GP methods arose from within the field of genetics, and they are built around classic assumptions about how genetic variation produces phenotypes. They fall into three broad families. Relationship-based models, such as ridge-regression BLUP and its equivalent GBLUP, model genetic relationships among individuals (VanRaden, 2008; Endelman, 2011). Marker-effect models, including the “Bayesian alphabet” of priors such as BayesA, BayesB and the Bayesian LASSO, encode assumptions about how marker effects are distributed (Meuwissen et al., 2001; de los Campos et al., 2013; Gianola, 2013). Kernel methods such as RKHS regression capture some non-additive genetic architectures (Gianola et al., 2006), and kernels built explicitly to model epistasis have been proposed (Jiang and Reif, 2015). Designing such models is a central activity of this field, and much of its literature consists of new priors, kernels or architectures, each tuned to the structure of genetic data.

General-purpose machine learning models have also been tested extensively for GP, with mixed results. They are sensitive to tuning and do not consistently beat linear models (Bellot et al., 2018; Pérez-Enciso and Zingaretti, 2019; Montesinos-López et al., 2021). The six-species benchmark of Azodi et al. (2019) compared 18 methods, from linear models to support vector regression, random forests, gradient boosting and neural networks, and found that no single method was best. Additionally, a recent ten-species benchmark found that tuned gradient-boosting and random-forest models were only marginally more accurate than Bayesian regressions (Quesada-Traver et al., 2025). Flexible machine learning models thus give small and inconsistent gains over methods that model genetics explicitly.

Sutton’s “Bitter Lesson” (Sutton, 2019) argues that, across AI research, general methods that leverage computation and learning have repeatedly overtaken methods built on human knowledge of the domain. In GP, the hand-designed methods above differ from one another by small, dataset-specific margins (Azodi et al., 2019; Quesada-Traver et al., 2025), and the Bitter Lesson predicts that larger gains will come instead from general learners that leverage computation. In most fields where the Bitter Lesson has played out, data and computation scale together, with larger models trained on ever larger datasets. In GP, however, datasets are often limited by a moderate number of available individuals who have been genotyped and phenotyped, so the computation and learning must happen elsewhere. TabPFN shifts that computation and learning to pretraining on millions of synthetic datasets, allowing it to learn general patterns of prediction that transfer to new problems without redesigning the model for each one.

TabPFN is a neural network method based on the transformer architecture that is designed to approximate Bayesian posterior prediction. As mentioned above, it comes pretrained on a large body of synthetic data. At inference, it takes the training rows of the focal dataset and their labels as context and predicts new rows in a single forward pass, without gradient or hyperparameter updates (Müller et al., 2022; Hollmann et al., 2023, 2025). The approach is analogous to a large language model, which is pretrained on a large body of text and when given a prompt as context, generates the continuation that best fits what it learned in pretraining. For TabPFN, the prompt is a table of labeled training rows and the continuation is a predictive distribution for each unlabeled row. TabPFN prediction has been strongest when *N* is on the order of hundreds to thousands, which is also the training-set size of many GP datasets. Until recently, however, its feature limits were far below *M* for a typical marker panel, which limited its utility in GP. Earlier TabPFN applications to GP therefore compressed or pre-selected features. As a recent example, in wheat, TabPFN served as one learner in an ensemble trained on SNPs selected with GWAS (Ren et al., 2025). The recent release of TabPFN-3.5 raised the feature limit to 20,000 columns (Prior Labs, 2026a,b). With these expanded feature limits, TabPFN-3.5 can be applied directly to larger marker panels and may improve on existing GP methods.

Here we take the Bitter Lesson approach to GP. We give TabPFN-3.5 the marker matrix directly, with no domain-specific modeling, no trait-based marker selection and no tuning, and ask whether a general-purpose pretrained model can match methods refined over two decades specifically for GP. We benchmark it on the 18 species-trait combinations of Azodi et al. (2019) and compare it with the 18 published methods. Because published results do not give their random train/test data splits, we also rerun GBLUP, RKHS and elastic-net models on identical splits to those given TabPFN-3.5. For the four species whose *M* exceeds TabPFN-3.5’s limit, we test a simple input of principal components plus evenly thinned markers. Because TabPFN-3.5 then sees only this hybrid input while the other models see every marker, we also fit GBLUP, RKHS and an elastic net to the same hybrid input, to separate the effect of the input from the effect of the model. Finally, we test whether TabPFN-3.5’s advantage holds in leave-families-out cross-validation, where models are trained on some soybean families and used to predict lines from families withheld from training. Among all 19 methods, TabPFN-3.5 was the most accurate on 11 of 18 tests and had the best median rank. On identical splits, it was significantly behind the best rerun model on only one test (spruce wood density) and ahead on 12 of 18, with significant gains on four: all three soybean traits and rice flowering time. When the other models were given the same hybrid input as TabPFN-3.5, it was significantly ahead on one test (rice flowering time) and behind on none. When whole soybean families were held out, however, TabPFN-3.5 lost its advantage and performed on par with GBLUP and RKHS.

## 2 Materials and methods

### 2.1 Data

We used the genotype, phenotype and fold-assignment files of Azodi et al. (2019), deposited on Dryad (doi:10.5061/dryad.xksn02vb9). The original studies that produced each dataset are cited in Table 1. Markers are coded −1/0/1, and the rice data also include a small fraction of imputed, non-integer dosages. Each species has three traits. The datasets span a 15-fold range in training-set size and a 58-fold range in marker number. Line counts in the deposit match those reported by Azodi et al. (2019), but three marker sets do not (maize 244,781 vs 332,178 reported; spruce 6,930 vs 6,932; soybean 4,234 vs 4,240).

**Table 1.** Datasets, from Azodi et al. (2019). Each species has three traits.

| Species | Population | Lines | Markers | Traits | Original data |
|---|---|---|---|---|---|
| Soybean | SoyNAM RILs | 5,014 | 4,234 | height, R8, yield | Xavier et al. (2016) |
| White spruce | tree breeding population | 1,722 | 6,930 | DBH, height, wood density | SmartForests project; SNP chip from Beaulieu et al. (2014) |
| Rice | IRRI elite breeding lines | 327 | 57,542 | flowering time, height, yield | Spindel et al. (2015) |
| Sorghum | diversity panel (NPGS) | 451 | 56,299 | grain moisture, height, yield | Fernandes et al. (2018) |
| Switchgrass | Northern Switchgrass Association Panel | 514 | 217,150 | anthesis date, height, standability | Lipka et al. (2014); Evans et al. (2015, 2018) |
| Maize | diversity panel | 391 | 244,781 | flowering time, height, yield | Hansey et al. (2011); Hirsch et al. (2014) |

### 2.2 Validation design

For each trait we used 10 replicate 80/20 holdouts, which matches the design of Azodi et al. (2019). Azodi et al. (2019) state that their training and testing designations are on Dryad, but the deposit holds 100 random five-fold partitions per species and does not say which defined their 10 test sets. We used fold 1 of the first 10 partitions (cv_1 to cv_10) as our test sets, so our splits follow their design but do not appear to be their splits. Accuracy is the Pearson correlation between predicted and observed phenotypes in the held-out lines, averaged over replicates, again matching Azodi et al. (2019). Within each split, every model was fit to the training lines only, so test-line phenotypes never entered model fitting. Test-line genotypes do enter the relationship matrices of the kernel models, which is standard for GBLUP and matches Azodi et al. (2019) (code at https://github.com/ShiuLab/Manuscript_Code).

#### Population-structure analyses (soybean)

To test generalization across families, we also built a leave-families-out design for soybean. In each of 10 replicates, 8 of the 38 SoyNAM families (21% of lines on average) were held out at random, and models were trained on the other 30 families. For each random split we divided accuracy into a between-family part (the correlation between observed and predicted family means of the test lines) and a within-family part (the correlation between deviations from those family means). We also fit GBLUP with family as a fixed effect, and TabPFN-3.5 with the 38 family labels added as one-hot columns given to every ensemble member.

### 2.3 TabPFN-3.5

We used `TabPFNRegressor` from the `tabpfn` Python package (version 9.1.0) with the TabPFN-3.5 weights (Prior Labs, 2026a), default settings, no hyperparameter tuning, and `random_state` set to the replicate number. TabPFN-3.5 does not make a single prediction per test line. Much like a random forest averages its trees, TabPFN-3.5 averages several forward passes of the model, called ensemble members, each run on a different subset of the input columns. In the tabpfn 9.1.0 package, each member sees at most 768 input columns. The default of 8 members can therefore include at most 8 × 768 = 6,144 columns between them, and with a wider input some markers would never be seen by any member. For these inputs we raised the number of members to the smallest value that covers every column at least once. For *M* input columns this is ⌈*M/*768⌉ members, so the 6,930 spruce markers needed ⌈6,930*/*768⌉ = 10. Soybean’s 4,234 markers fit within the default 8 members.

#### Hybrid input (rice, sorghum, maize, switchgrass)

These species have about 56,000–245,000 markers, more than the 20,000 columns TabPFN-3.5 accepts (Prior Labs, 2026b). For each replicate we supplied the top 100 principal components of all markers, computed on the training lines only with test lines projected onto them, and 19,900 raw markers chosen at evenly spaced ranks after sorting by chromosome and position. We used the `tabpfn` package’s `constant_and_balanced` feature subsampling so that the 100 principal components are given to every ensemble member and the thinned markers are divided among members. Each member therefore sees all 100 components plus 668 of the markers, and 30 members are needed to include all 19,900 markers.

#### Alternative inputs

We also ran all principal components alone on the four species with more than 20,000 markers. These panels have at most 411 components, fewer than the 768 columns each ensemble member sees, so every member saw all of them. We also ran thinned markers alone on the same four species and, for rice, all 57,542 markers with the package’s pretraining limits disabled (`ignore_pretraining_limits`).

### 2.4 Comparison models

All comparison models were fit on exactly the same training and test lines as TabPFN-3.5. They include:

- **GBLUP**. *y* = *µ* + *g* + *e* with 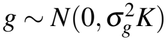, fit by REML on the training lines. Predictions are the BLUPs of *g* for test lines. *K* was built as in the rrBLUP script of Azodi et al.: 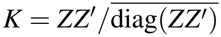, with *Z* the column-standardized marker matrix of all lines.
- **RKHS**. The same REML model with a Gaussian kernel 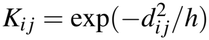, where 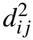 is the mean squared difference between standardized marker profiles and *h* is its median over pairs (Gianola et al., 2006; de los Campos et al., 2013).
- **Elastic net (ENet)**. Penalized regression with an L1/L2 mixing ratio of 0.5 and the penalty chosen from 30 values by 5-fold cross-validation within the training lines (scikit-learn `ElasticNetCV`; Pedregosa et al., 2011). Before fitting, every input column was standardized to mean zero and unit variance using the training lines. We call this elastic net “ENet” to distinguish it from the EN5 and EN11 ensembles of Azodi et al. (2019).

For the four species with more than 20,000 markers, GBLUP, RKHS and ENet were each fit twice, once on all markers and once on the same 100 PCs + 19,900 thinned markers that TabPFN-3.5 received (GBLUP-hybrid, RKHS-hybrid and ENet-hybrid).

### 2.5 Comparison with published results

Table S5 of Azodi et al. (2019) gives the mean accuracy of 18 methods for each species and trait: rrBLUP, Bayesian ridge regression, BayesA, BayesB, Bayesian LASSO, three support vector regressions, random forest, gradient tree boosting, five neural network variants, a convolutional network, and two ensembles (EN5, EN11) that average 5 or 11 of these methods. We ranked TabPFN-3.5’s mean accuracy among these 18 published means. Because published and TabPFN-3.5 means come from different data splits, this ranking is descriptive. The comparison models rerun on identical splits carry the formal comparisons.

### 2.6 Statistics

For each species and trait we computed the per-replicate difference in accuracy between TabPFN-3.5 and each comparison model. Within each replicate the training and test lines are disjoint, but across replicates both training sets and test sets overlap (any two test sets share about 20% of their lines), so a naive paired t-test that treats replicates as independent understates the variance. We therefore report the mean difference with a 95% interval from the corrected resampled t-test of Nadeau and Bengio (2003), using a test-to-training size ratio of 20/80 = 0.25, along with the number of replicates in which TabPFN-3.5 was more accurate.

### 2.7 Software and compute

TabPFN-3.5 was run on the integrated GPU of an Apple M4 Pro chip. Comparison models ran on CPU, either on a Linux cloud machine or the same Apple M4 Pro laptop, in Python 3.13 and scikit-learn 1.9.1. Run times are reported per fit.

## 3 Results

### 3.1 Against published methods, TabPFN-3.5 has the best median rank

With no tuning, and among 19 methods (itself plus the 18 published), TabPFN-3.5 had the highest mean accuracy on 11 of 18 species-trait tests and was in the top three on one more. The next best method, the EN11 ensemble of 11 methods from Azodi et al. (2019), had a median rank of 3.5 (Table 2).

**Table 2.** Rank of TabPFN-3.5 among the 18 published GP results in Azodi et al. (2019) (Table S5 of that paper), over the 18 species-trait tests. Only the seven methods with the best median ranks are shown.

| Method | Median rank (of 19) | Tests ranked first | Tests in top 3 |
|---|---|---|---|
| TabPFN-3.5 | 1 | 11 | 12 |
| EN11 (ensemble of 11 methods) | 3.5 | 2 | 9 |
| BayesA | 5 | 0 | 7 |
| EN5 (ensemble of 5 methods) | 5.5 | 0 | 4 |
| Bayesian ridge regression | 6 | 2 | 2 |
| rrBLUP | 6 | 0 | 6 |
| BayesB | 7 | 2 | 2 |

On the six tests where it saw every marker (soybean, spruce), TabPFN-3.5 ranked first every time. On the hybrid input (rice, sorghum, maize, switchgrass), it ranked first on 5 of 12. Its weakest species was sorghum, where it ranked 8th to 13th. Per-test accuracies for every model are in Table S1.

These ranks compare TabPFN-3.5 with published means from different random data splits, and that difference appears to matter. On the five species other than rice, our GBLUP (equivalent to rrBLUP) was within about 0.04 in accuracy (Pearson’s r) of the published rrBLUP on every trait. On rice it was far higher: 0.186 higher for yield, 0.140 higher for flowering time, and 0.053 higher for height. The published rice rrBLUP appears anomalous. In Azodi et al.’s own results, rrBLUP and Bayesian ridge regression, which fit similar models, agree within 0.02 on all 15 non-rice tests but differ by 0.04–0.12 on rice. Against their Bayesian ridge regression, our GBLUP agrees more closely for rice flowering time (0.557 vs 0.539) and height (0.396 vs 0.385) but remains higher for yield (0.458 vs 0.363), a gap we cannot explain. As a check, our own GBLUP, rerun on our splits, also ranked first on 5 of 18 tests against the published table, so part of TabPFN-3.5’s lead in this ranking may reflect the splits rather than the model. Below is a comparison of a smaller number of models on identical splits.

### 3.2 On identical splits, TabPFN-3.5 matches or beats strong comparison models, with clear wins in soybean

Compared with the most accurate of GBLUP, RKHS and ENet on the same splits, each given either all markers or the hybrid input, TabPFN-3.5’s mean accuracy was higher on 12 of 18 tests (Figure 1). TabPFN-3.5 was significantly better on four tests (blue in Figure 1), all three soybean traits (+0.017 to +0.041) and rice flowering time (+0.047). It was significantly worse on one (red), spruce wood density (−0.032). On the other 13 tests (grey) the interval included zero, so those differences cannot be distinguished from split-to-split noise.

**Figure 1:**
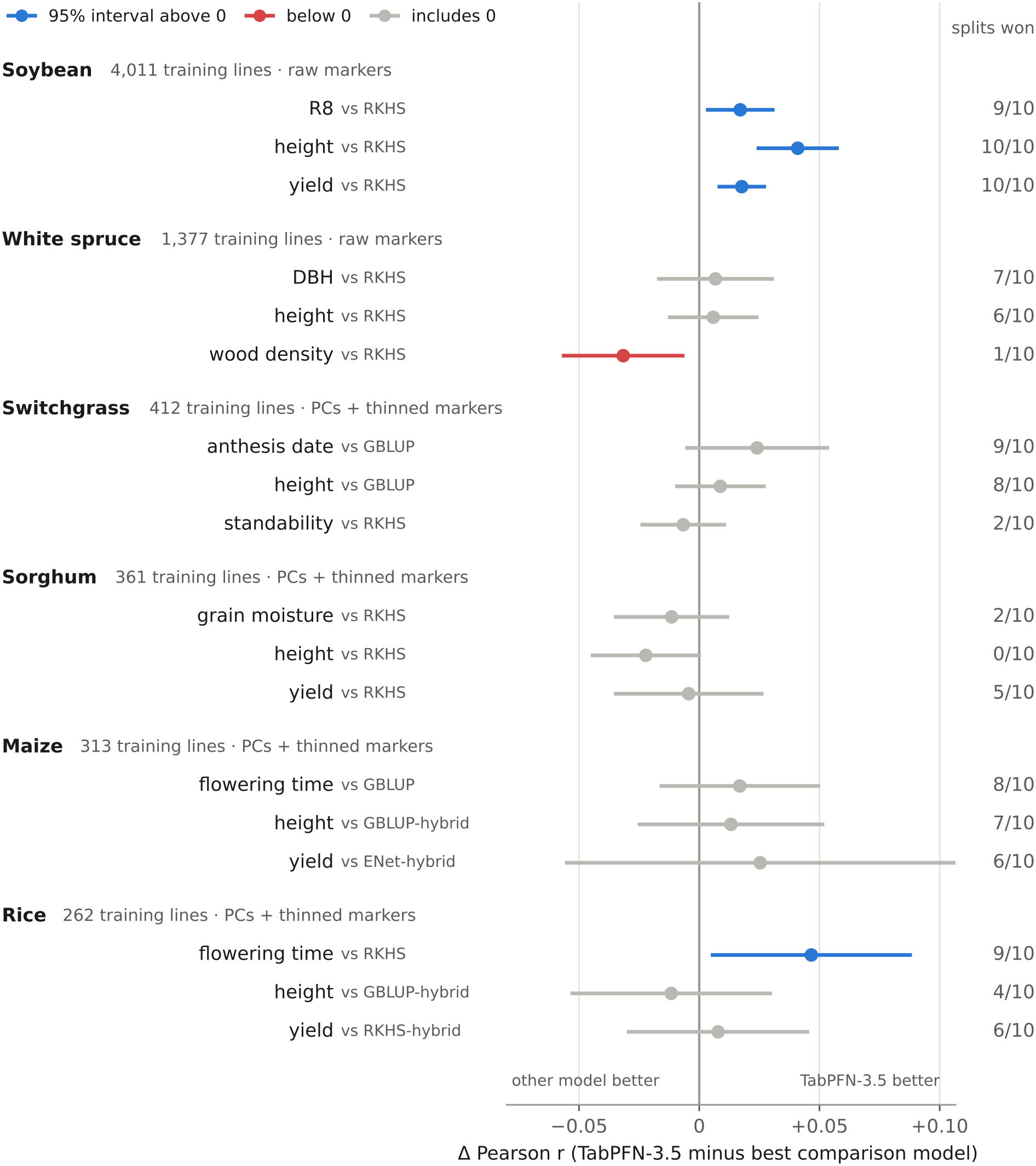
TabPFN-3.5 matches or beats the comparison models on identical splits, with clear gains in soybean. Each point is the mean difference in Pearson *r* between TabPFN-3.5 and the most accurate comparison model for that test (GBLUP, RKHS or ENet; on all markers or on the hybrid input), over the same 10 replicate 80/20 splits. Horizontal bars are 95% intervals from the corrected resampled t-test (Nadeau and Bengio, 2003). Blue: interval above zero; red: below zero; grey: includes zero. The right-hand column gives the number of splits on which TabPFN-3.5 was more accurate. Species are ordered by training-set size, and the label beside each gives TabPFN-3.5’s input: all raw markers, or 100 principal components plus 19,900 thinned markers.

Soybean is the one species where TabPFN-3.5 won every trait. It is also the dataset with the most lines which results in tighter confidence intervals. These comparisons are also conservative for TabPFN-3.5 because of the winner’s curse phenomenon. For each test, TabPFN-3.5 is compared with whichever comparison model and input scored highest on these same splits, and that best score tends to overstate the chosen model’s true accuracy.

Among the comparison models (i.e. excluding TabPFN-3.5), RKHS was strongest on 12 of 18 tests. It was not among the methods of Azodi et al. (2019), yet on spruce it matched or beat every published method. ENet was strongest on only one test, maize yield, where on the hybrid input it reached 0.560 against 0.543 for GBLUP-hybrid. ENet’s accuracy there varied widely between splits (*r* = 0.48 to 0.67), so TabPFN-3.5’s lead over it (+0.025) fell within split-to-split noise, whereas its lead over GBLUP-hybrid for maize yield (+0.042) was significant. On every other test ENet trailed GBLUP, and it trailed TabPFN-3.5 on every trait.

### 3.3 The hybrid input works, and reducing markers costs GBLUP and RKHS almost nothing

Before comparing TabPFN-3.5 with other models, we asked how its own accuracy depends on its input. For the four species with more markers than it accepts, we gave TabPFN-3.5 three inputs, all principal components alone, 19,900 thinned markers alone, and the hybrid of the two. The hybrid input was the most accurate on 9 of 12 traits (Table 3). Thinned markers alone did better on rice flowering time and yield, and principal components alone on sorghum grain moisture. Across the 120 species-by-trait-by-split comparisons, the hybrid input beat thinned markers alone on 97 splits and principal components alone on 82 splits. For GBLUP and RKHS, switching from all markers to the hybrid input changed accuracy by at most 0.008 on any trait. The elastic net was more sensitive. The hybrid input suited it better on rice yield (+0.093) and maize yield (+0.064), and worse on rice height (−0.075).

**Table 3.** Accuracy (Pearson *r*) of each model on each input, by trait, for the four species with more markers than TabPFN-3.5 accepts. “PCs” is all principal components, “all” is all markers, and “hybrid” is 100 PCs + 19,900 thinned markers. TabPFN-3.5 on all markers was run for rice only, with `ignore_pretraining_limits`. Traits are AN (anthesis date), FT (flowering time), HT (height), MO (grain moisture), ST (standability) and YLD (yield). Values are means over 10 splits. The best value for each trait is in bold, including ties. *In 4 of 10 rice yield fits on all markers and 2 of 10 on the hybrid input, the elastic net kept no markers and predicted a constant, so the correlation is undefined. Those fits are counted as *r* = 0.

|  | Switchgrass |  |  | Sorghum |  |  | Maize |  |  | Rice |  |  |
|---|---|---|---|---|---|---|---|---|---|---|---|---|
| Model and input | AN | HT | ST | MO | HT | YLD | FT | HT | YLD | FT | HT | YLD |
| TabPFN-3.5, PCs | 0.889 | 0.618 | 0.675 | 0.535 | 0.597 | 0.382 | 0.725 | 0.497 | 0.584 | 0.564 | 0.378 | 0.407 |
| TabPFN-3.5, thinned | 0.868 | 0.598 | 0.667 | 0.513 | 0.579 | 0.375 | 0.715 | 0.489 | 0.553 | **0.617** | 0.367 | **0.482** |
| TabPFN-3.5, hybrid | **0.897** | **0.620** | 0.685 | 0.527 | 0.600 | 0.397 | **0.741** | **0.502** | **0.585** | 0.612 | 0.388 | 0.471 |
| TabPFN-3.5, all | – | – | – | – | – | – | – | – | – | 0.603 | 0.372 | **0.482** |
| GBLUP, all | 0.873 | 0.611 | **0.691** | 0.536 | 0.619 | 0.393 | 0.724 | 0.488 | 0.536 | 0.557 | 0.396 | 0.458 |
| GBLUP, hybrid | 0.870 | 0.606 | 0.683 | 0.530 | 0.616 | 0.391 | 0.717 | 0.489 | 0.543 | 0.549 | **0.400** | 0.460 |
| RKHS, all | 0.870 | 0.611 | **0.691** | **0.539** | **0.622** | **0.402** | 0.715 | 0.475 | 0.529 | 0.566 | 0.383 | 0.460 |
| RKHS, hybrid | 0.868 | 0.607 | 0.686 | 0.534 | 0.620 | 0.401 | 0.710 | 0.477 | 0.535 | 0.558 | 0.382 | 0.463 |
| ENet, all | 0.829 | 0.566 | 0.657 | 0.496 | 0.548 | 0.300 | 0.692 | 0.408 | 0.496 | 0.504 | 0.355 | 0.218* |
| ENet, hybrid | 0.832 | 0.561 | 0.631 | 0.489 | 0.559 | 0.312 | 0.667 | 0.411 | 0.560 | 0.480 | 0.280 | 0.311* |

The 20,000-feature limit is not absolute. The `ignore_pretraining_limits` option lets TabPFN-3.5 run on wider inputs, with a warning that the input lies outside the range the model was pretrained on and that accuracy may suffer. Using this option, we tested TabPFN-3.5 on all 57,542 rice markers. For flowering time, height and yield, all markers gave *r* = 0.603, 0.372 and 0.482, against 0.617, 0.367 and 0.482 on thinned markers and 0.612, 0.388 and 0.471 on the hybrid input. All markers beat the hybrid input on only 12 of 30 trait-by-split comparisons and took nearly three times as long per fit. So the hybrid input remained the better choice here, but the `ignore_pretraining_limits` option is available for panels where thinning might discard more information.

Giving every model the same hybrid input asks a second question, whether TabPFN-3.5’s advantage comes from the input rather than the model. Against the most accurate of GBLUP-hybrid, RKHS-hybrid and ENet-hybrid, TabPFN-3.5 was ahead on 7 of the 12 tests in the four species with more than 20,000 markers, significantly so on rice flowering time (+0.055), and significantly behind on none. Thus we conclude that the hybrid input itself is not an advantage. Where TabPFN-3.5 leads, the lead appears to come from the model, not the hybrid input.

### 3.4 On held-out soybean families, TabPFN-3.5 performs on par with GBLUP and RKHS

Random splits in the soybean SoyNAM families put sibling lines in both training and test sets, which inflates accuracy in structured populations (Habier et al., 2007; Werner et al., 2020). When whole SoyNAM families were held out, accuracy fell sharply for every model, TabPFN-3.5 included (Table 4).

**Table 4.**
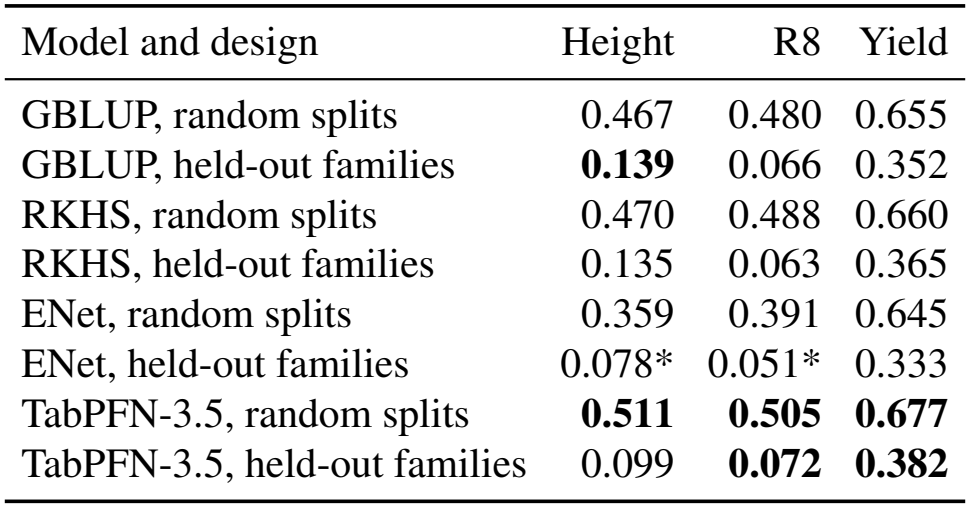
Soybean accuracy (Pearson *r*) on random 80/20 splits and with whole SoyNAM families held out, for each model. The best value for each trait within each design is in bold. *In 4 of 10 height fits and 2 of 10 R8 fits, the elastic net kept no markers and predicted a constant, so the correlation is undefined. Those fits are counted as *r* = 0.

| Model and design | Height | R8 | Yield |
|---|---|---|---|
| GBLUP, random splits | 0.467 | 0.480 | 0.655 |
| GBLUP, held-out families | **0.139** | 0.066 | 0.352 |
| RKHS, random splits | 0.470 | 0.488 | 0.660 |
| RKHS, held-out families | 0.135 | 0.063 | 0.365 |
| ENet, random splits | 0.359 | 0.391 | 0.645 |
| ENet, held-out families | 0.078* | 0.051* | 0.333 |
| TabPFN-3.5, random splits | **0.511** | **0.505** | **0.677** |
| TabPFN-3.5, held-out families | 0.099 | **0.072** | **0.382** |

Accuracy depended heavily on which eight families were held out (for yield, r ranged from 0.10 to 0.59 across splits), so TabPFN-3.5’s differences from GBLUP and RKHS were indistinguishable from zero on all three traits, and the only significant difference was a small yield lead over the elastic net.

These results suggest that TabPFN-3.5’s advantage in soybean depends on the genetic backgrounds represented in the training population. When entire SoyNAM families were held out, its advantage over other models largely disappeared. This pattern does not indicate a limitation unique to foundation models like TabPFN. Rather, it highlights a broader challenge for genomic prediction when relationships between training and test individuals are reduced. TabPFN-3.5 can compete with and beat specialized methods when predicting within the sampled population, but in soybean these results provide no evidence that it generalizes better or worse to families absent from training.

### 3.5 Run time

Each TabPFN-3.5 fit took 22–93 s on the integrated GPU of an Apple M4 Pro, against seconds for GBLUP and RKHS and less than a few minutes for ENet on CPU. TabPFN-3.5 needs no tuning, while some of the published methods required grid searches with repeated cross-validation (Azodi et al., 2019). Thus even on a modest laptop GPU, TabPFN-3.5 genomic prediction runs are not appreciably slower than other CPU-based methods.

## 4 Discussion

Given only the marker matrix and no tuning, TabPFN-3.5 was ahead of the strongest GP model on 12 of 18 published tests and significantly behind on just one. It was clearly worse only on spruce wood density, and clearly better on all three soybean traits and on rice flowering time, where its lead held when the other models were given the same hybrid input. When the test lines came from soybean families absent from training, it performed on par with GBLUP and RKHS.

The gains are modest in absolute terms: +0.017 to +0.041 in correlation on soybean, or 3–9% relative to the best comparison model. They are similar in size to the differences that separate established GP methods in past studies (Azodi et al., 2019; Quesada-Traver et al., 2025).

### 4.1 Where TabPFN-3.5 wins

TabPFN-3.5’s advantage was clearest in soybean, the dataset with the most training lines, where differences are measured most precisely. With only 262–412 training lines in the four small panels (rice, maize, sorghum and switchgrass), a real accuracy advantage of ~0.02 would often go undetected. Training-set size is confounded here with population type, however, because soybean is a set of related biparental families while the small panels are diverse collections. TabPFN-3.5’s loss of advantage when whole families were held out points to population structure as the driver of its advantage when soybean families are split between training and test sets.

A follow-up analysis adds detail. On random splits in soybean, most of every model’s accuracy came from predicting differences between families (between-family *r* from 0.92 to 0.98), while accuracy among sibs within a family was low (*r* from 0.03 to 0.19). TabPFN-3.5 ranked families no better than GBLUP and RKHS did, and ranked sibs within a family no better on height and worse on R8 and yield. Its advantage came from weighting. It put more of its predicted variation on family means (86 to 94%, against 69 to 84% for GBLUP and RKHS), whose differences are large and well estimated. And it gave correspondingly less weight to the noisier differences within families. To check this inference we explicitly gave GBLUP and TabPFN-3.5 each line’s family (a fixed effect in GBLUP, one-hot columns in TabPFN-3.5) and refit them. With family as a fixed effect, GBLUP rose to *r* = 0.519, 0.532 and 0.699 for height, R8 and yield, and TabPFN-3.5 given the same labels reached 0.533, 0.530 and 0.696. Thus, from the markers alone, TabPFN-3.5 recovered roughly half or more of the accuracy gain that explicit family labels provide. Because this advantage depends on having full sibs from the same family in training, it does not carry over to new families.

These results hint at two strengths of a model that learns how to predict from the data, compared with GBLUP, whose form (additive marker effects, a fixed relationship matrix and a single variance component) is set in advance. GBLUP applies a single variance component, and so a single degree of shrinkage, to both family means and the differences within families, whereas TabPFN-3.5 shrank the two differently without being told that families existed. It also appeared to partially recover the family structure itself despite not being given it as an input. Breeding populations may carry similar unprovided or perhaps unknown structures, and a model that learns them from the data rather than requiring them to be specified may be an advantage in practice. Whether this holds where the structure is subtler than SoyNAM’s 38 biparental families remains untested.

Having full-sib relatives in the training and test sets is not uncommon in breeding. In plant breeding, sparse testing and early-generation selection frequently involve full sibs or other first-degree relatives. In animal breeding, pigs, poultry and especially aquaculture species such as salmon, tilapia and shrimp are often bred in large full-sib families, and traits are routinely measured on sibs of the selection candidates. The datasets we analyzed lack the multi-cycle structure needed to test whether TabPFN helps when a model trained on one breeding cycle predicts new crosses in the next, and this would be an interesting avenue for future research.

### 4.2 A test of the Bitter Lesson in genomic prediction

Our results are consistent with the Bitter Lesson in GP. TabPFN-3.5, a model whose only knowledge comes from pretraining on synthetic data, reached the accuracy of methods refined over two decades for this problem, and exceeded it on random splits where data were most plentiful. The test is not entirely pure. For the four species with more than 20,000 markers, the feature limit forced us to reduce the marker input which imports some human design and taste in choosing principal components plus even marker thinning. That said, the knowledge involved is fairly generic. Principal component analysis is a general-purpose data-reduction method, and thinning relies only on marker order and the expectation that nearby markers are correlated through linkage. Both make much weaker assumptions about how genotype maps to phenotype than do relationship matrices, marker-effect priors and kernels of other GP methods. Furthermore, future TabPFN releases that accept more features could allow full marker panels for more of these datasets, removing even this step, and these comparisons are worth repeating when they arrive.

TabPFN leverages computation uniquely compared to the other models in this study. Other methods fit their parameters to each training set from scratch, so once the model is specified, their accuracy is bounded by the data and by the assumptions built into the model. TabPFN does not fit parameters to the training lines at all. It reads them as context and predicts the test lines in a forward pass, a process known as in-context learning (Müller et al., 2022; Hollmann et al., 2025). What it knows about prediction was learned earlier, when it was pretrained on large synthetic datasets. This pretraining can scale as computation grows by using larger models, more synthetic datasets, and longer training. So a model of this kind could improve on GP problems even when the GP data stay the same. In-context learning therefore offers GP a path to higher accuracy that grows with computation rather than with expert knowledge, which is the path the Bitter Lesson predicts will one day win out.

## Use of generative AI

Claude (Anthropic) was used to write and run analysis code for the comparison models, to verify citations, and to draft and revise the manuscript text. ChatGPT (OpenAI) was used to review a draft. The author designed the study, ran the analyses, checked all results, edited the text, and takes full responsibility for the content.

## Data and code availability

All data are from Azodi et al. (2019), deposited on Dryad under CC0 (doi:10.5061/dryad.xksn02vb9), and a checksum-verified mirror is included in the code repository. Code, per-replicate results for every model, and scripts to regenerate all tables and figures are at github.com/flag0010/tabpfn-gp-manuscript (MIT License). The TabPFN-3.5 weights are licensed separately by Prior Labs under a non-commercial license.

## Supplementary material

**Table S1:** Mean accuracy (Pearson *r*) of every model on each species-trait test. TabPFN-3.5 and the comparison models are means over the same 10 replicate 80/20 splits. TabPFN-3.5 received raw markers for soybean and spruce and the hybrid input (100 PCs + 19,900 thinned markers) for the other four species. GBLUP, RKHS and ENet used all markers. For the four species with more than 20,000 markers, GBLUP-hybrid, RKHS-hybrid and ENet-hybrid were also fit to the same 100 PCs + 19,900 thinned markers that TabPFN-3.5 received. Fits where ENet kept no markers are counted as *r* = 0. Best published values are from Table S5 of Azodi et al. (2019), computed on different random splits. The last column ranks TabPFN-3.5 among those 18 published means.

| Species | Trait | TabPFN-3.5 | GBLUP | RKHS | ENet | GBLUP-hybrid | RKHS-hybrid | ENet-hybrid | Best published (method) | TabPFN-3.5 rank of 19 |
|---|---|---|---|---|---|---|---|---|---|---|
| Soybean | R8 | 0.505 | 0.480 | 0.488 | 0.391 | – | – | – | 0.492 (EN11) | 1 |
| Soybean | height | 0.511 | 0.467 | 0.470 | 0.359 | – | – | – | 0.468 (BayesA) | 1 |
| Soybean | yield | 0.677 | 0.655 | 0.660 | 0.645 | – | – | – | 0.650 (EN11) | 1 |
| White spruce | DBH | 0.279 | 0.253 | 0.272 | 0.225 | – | – | – | 0.275 (RF) | 1 |
| White spruce | height | 0.350 | 0.328 | 0.344 | 0.299 | – | – | – | 0.346 (RF) | 1 |
| White spruce | wood density | 0.405 | 0.429 | 0.437 | 0.395 | – | – | – | 0.398 (EN11) | 1 |
| Switchgrass | anthesis date | 0.897 | 0.873 | 0.870 | 0.829 | 0.870 | 0.868 | 0.832 | 0.855 (rrBLUP) | 1 |
| Switchgrass | height | 0.620 | 0.611 | 0.611 | 0.566 | 0.606 | 0.607 | 0.561 | 0.615 (SVRpoly) | 1 |
| Switchgrass | standability | 0.685 | 0.691 | 0.691 | 0.657 | 0.683 | 0.686 | 0.631 | 0.693 (SVRrbf) | 9 |
| Sorghum | grain moisture | 0.527 | 0.536 | 0.539 | 0.496 | 0.530 | 0.534 | 0.489 | 0.572 (EN11) | 13 |
| Sorghum | height | 0.600 | 0.619 | 0.622 | 0.548 | 0.616 | 0.620 | 0.559 | 0.628 (BayesB) | 11 |
| Sorghum | yield | 0.397 | 0.393 | 0.402 | 0.300 | 0.391 | 0.401 | 0.312 | 0.411 (BRR) | 8 |
| Maize | flowering time | 0.741 | 0.724 | 0.715 | 0.692 | 0.717 | 0.710 | 0.667 | 0.765 (BRR) | 8 |
| Maize | height | 0.502 | 0.488 | 0.475 | 0.408 | 0.489 | 0.477 | 0.411 | 0.448 (EN5) | 1 |
| Maize | yield | 0.585 | 0.536 | 0.529 | 0.496 | 0.543 | 0.535 | 0.560 | 0.546 (EN5) | 1 |
| Rice | flowering time | 0.612 | 0.557 | 0.566 | 0.504 | 0.549 | 0.558 | 0.480 | 0.650 (BayesB) | 3 |
| Rice | height | 0.388 | 0.396 | 0.383 | 0.355 | 0.400 | 0.382 | 0.280 | 0.408 (EN11) | 5 |
| Rice | yield | 0.471 | 0.458 | 0.460 | 0.218 | 0.460 | 0.463 | 0.311 | 0.385 (EN11) | 1 |

